# *Azotobacter vinelandii* glutaredoxin D delivers the core [Fe_2_S_2_] cluster to nitrogenase cofactor scaffold protein NifU

**DOI:** 10.1101/2025.10.21.683637

**Authors:** Juan Andrés Collantes-García, Elena Rosa-Núñez, Alejandro M Armas, Daniel Raimunda, Ana Pérez-González, Yisong Guo, Carlos Echávarri-Erasun, Luis M. Rubio, Manuel González-Guerrero

## Abstract

The scaffold protein NifU plays a central role in assembling the precursor [Fe_4_S_4_] clusters required for nitrogenase to function. The synthesis of these precursors depends on a catalytic [Fe_2_S_2_] group within NifU core ferredoxin domain. Here, we show that the monothiol glutaredoxin GrxD delivers this cluster to the NifU scaffold protein. Consistently, *grxD* mutants have reduced nitrogenase activity, the result of altered iron allocation to this enzyme. Biochemical assays show that GrxD unidirectionally transfers [Fe_2_S_2_] to NifU through protein-protein interaction. This allows GrxD to restore apo-NifU functionality, enabling proper [Fe_4_S_4_] synthesis, and NifH activation. These findings are crucial to understand how iron is allocated to nitrogenase for biological nitrogen fixation.

## Introduction

Biological nitrogen fixation is carried out by the metalloenzyme nitrogenase (1). The molybdenum nitrogenases are constituted by three Nif (nitrogen fixation) proteins, NifH, NifD, and NifK organized in two Components (2). Component I is a NifD_2_K_2_ heterotetramer in which the catalysis takes place. Component II is a NifH dimer that provides electrons to NifDK for the catalysis. The complex stoichiometry is two NifH dimers per NifDK tetramer. The molybdenum nitrogenases require three different iron-sulfur clusters to operate: two [Fe_4_S_4_], two P-clusters and two iron-molybdenum cofactors (FeMo-co) (3, 4). The [Fe_4_S_4_] clusters are inserted in each of the two NifH dimers, specifically at the interface of the subunits. Component I contains the P-clusters ([Fe_8_S_7_]) at the interfaces of each of the two NifDK halves, and a FeMoco, [Fe_7_S_9_CMo-*R*-homocitrate]) at the active site of each NifD subunit. This arrangement of cofactors allows for electrons to be channeled from the [Fe_4_S_4_] cluster to FeMo-co through the P-cluster (5, 6).

The biosynthesis of the nitrogenase clusters is a complex process, involving several additional Nif proteins (7). It is initiated by the assembly of transient [Fe_4_S_4_] clusters on the scaffold protein NifU. This is a 33 kDa dimer with each monomer comprised of three different domains: an IscU-like N-terminal domain, a core ferredoxin domain with a permanent catalytical [Fe_2_S_2_] cluster, and a C-terminal Nfu-like region (8, 9). The transient clusters are assembled on the N-terminal domain initially as [Fe_2_S_2_] groups. Sulfur is provided by cysteine desulfurase NifS, while the iron donor is yet-to-be determined (10). The synthesized clusters are condensed into [Fe_4_S_4_] and moved to the C-domain. Subsequently, these groups are transferred to NifH, to NifB to produce the NifB-cofactor precursor of FeMoco, and to NifQ to integrate Mo into FeMo-co synthesis (8, 11, 12), among other candidate acceptor proteins.

In addition to the iron-donor for the synthesis of the transient [Fe_2_S_2_] groups, NifU requires a second iron source to donate the catalytic [Fe_2_S_2_] cluster. This group must be synthesized by a “housekeeping” scaffold protein, responsible for providing the [FeS] clusters to enzymes not involved in diazotrophy, since the *nif* genes are repressed when sufficient fixed nitrogen is available (13). In model diazotroph *Azotobacter vinelandii*, IscU would be the source of these groups, as it is the one scaffold protein produced under non-diazotrophic conditions (14). From this protein, the clusters will be transferred to carrier proteins that will dock with specific acceptors and deliver the metal clusters to them. The monothiol glutaredoxin GrxD (Avin 14040, also known as Grx5) is one of these carriers able to accept a [Fe_2_S_2_] cluster from IscU (15). This class of glutaredoxins typically delivers a [Fe_2_S_2_] group coordinated by a Grx homodimer, or a Grx-BolA heterodimer (16, 17). Interestingly, *grxD* is among the first up-regulated genes as *A. vinelandii* transitions into a diazotrophic metabolism (18, 19). This expression profile indicates a role in early delivery of [Fe_2_S_2_] clusters to nascent [FeS] proteins involved in biological nitrogen fixation. Consequently, NifU could be among the GrxD acceptors, as it plays a central role in nitrogenase cofactor synthesis, and it requires a catalytic [Fe_2_S_2_] cluster to operate (8, 20). Supporting this hypothesis, here we report how optimal nitrogenase activity relies on *grxD* expression, and how GrxD transfer of a [Fe_2_S_2_] cluster to the core domain of NifU is essential for NifU activity.

## Results

### *GrxD is required for optimal nitrogen fixation in* Azotobacter vinelandii

To determine the role of GrxD in biological nitrogen fixation, an in-frame deletion was generated in the *Avin14040* gene of the *A. vinelandii* DJ (wild-type) chromosome. The resulting strain, DJ3045 (Table 1), presented slower diazotrophic growth than the DJ strain (Fig. 1A). This mutation also resulted in DJ3045 cells having 50 % less iron than the DJ strain (Fig. 1B). In contrast, no significant differences were observed when fixed nitrogen (ammonium) was present in the medium (Fig. 1C), even with low iron in the growth medium (Fig. S1). Consistent with a role of GrxD in diazotrophy, DJ3045 had lower nitrogenase activity than the controls (Fig. 1D). The diazotrophic growth, iron content and nitrogenase activity were restored when a wild-type copy of *grxD* was reintroduced into DJ3045 (strain DC47, Figure 1 and Table 1).

**Table 1.** *Azotobacter vinelandii* strains use in this work.

| Strain | Genotype | Source |
| --- | --- | --- |
| DJ | Highly transformable variant of OP | 49 |
| DJ3045 | $\Delta grxD$ mutant in frame from amino acid 2 to 50 | This work |
| DC47 | GrxD complementation | This work |
| DJ33 | $\Delta nifDK$ | 47 |

**Figure 1.**
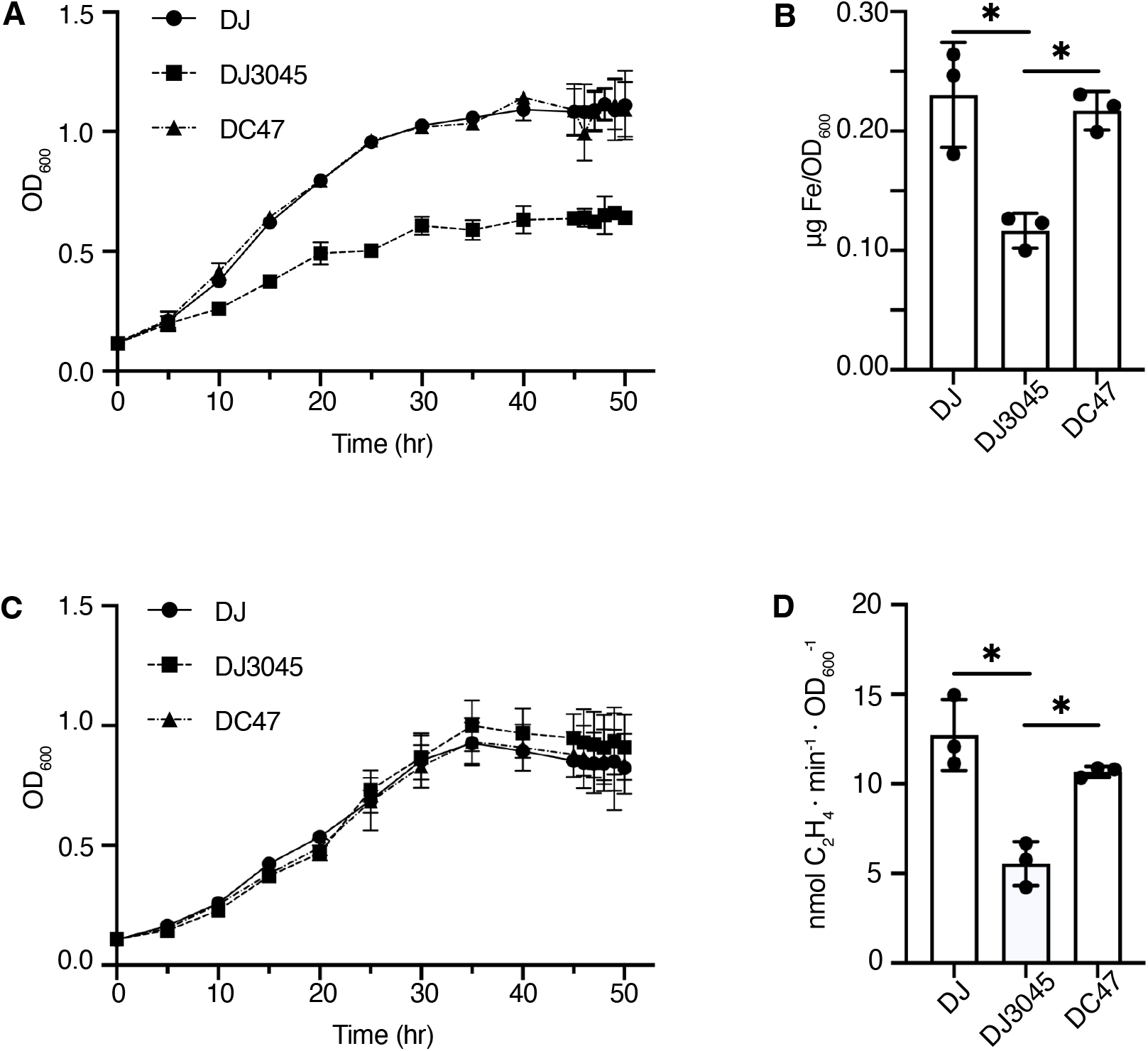
Mutation in *grxD* alters diazotrophic growth. A. Growth under diazotrophic conditions of wild type *A. vinelandii* strain (DJ), *grxD* in-frame mutant (DJ3045), and DJ3045 transformed with a wild-type copy of *grxD* (DC47). B. Iron content of DJ, DJ3045, and DC47 strains grown in diazotrophic conditions. C. Growth under non-diazotrophic conditions of wild type DJ, DJ3045, and DC47. D. Nitrogenase activity assay of DJ, DJ3045, and DC47 cells 4 h after de-repression. Bars represent the average ± SD (n=3).

### GrxD interacts with NifU

The phenotype of DJ3045 suggested impaired iron metabolism during diazotrophic growth. Considering the important role that NifU plays in iron allocation to Nif proteins and its reliance on its catalytic [Fe_2_S_2_] cluster, such as those provided by monothiol glutaredoxins (8, 21), the direct interaction between these two proteins was tested. To this end, a C-terminal Strep-tagged *A. vinelandii* GrxD (GrxD_S_) was produced and purified with a 1.25:1 iron:GrxD ratio (Table 2), indicative of a [Fe_2_S_2_] cluster coordinated by two GrxDs. This protein was incubated with a purified N-terminal His-tagged *A. vinelandii* NifU (_H_NifU), containing the two iron per monomer characteristic of the core [Fe_2_S_2_] (Table 2), under anaerobic conditions and subsequently loaded onto a Strep-tactin column. In the elutions, _H_NifU was detected together with GrxD_S_ (Fig. 2, Fig. S2). _H_NifU presence in the elution fractions was dependent of interaction with GrxD_S_; when GrxD_S_ was not present, no NifU could be detected in the elutions (Figure S3). Similarly, N-terminal His-tagged GrxD could be co-purified with a C-terminal Strep-tagged NifU (Fig. S4, Table 2).

**Table 2.**
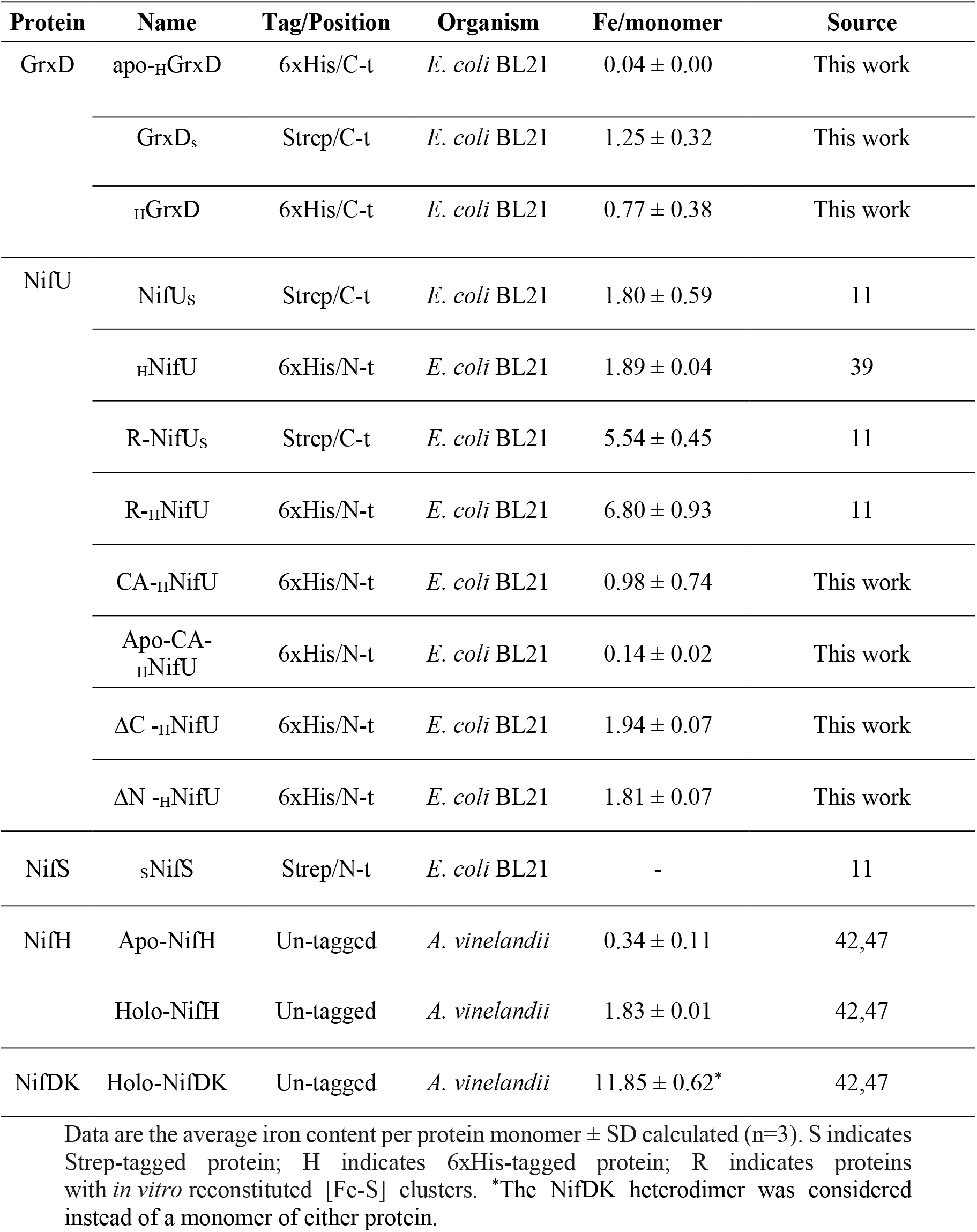
Proteins used in this study.

| Protein | Name | Tag/Position | Organism | Fe/monomer | Source |
| --- | --- | --- | --- | --- | --- |
| GrxD | apo-HGrxD | 6xHis/C-t | *E. coli* BL21 | $0.04 \pm 0.00$ | This work |
| | GrxD<sub>s</sub> | Strep/C-t | *E. coli* BL21 | $1.25 \pm 0.32$ | This work |
| | HGrxD | 6xHis/C-t | *E. coli* BL21 | $0.77 \pm 0.38$ | This work |
| NifU | NifU<sub>s</sub> | Strep/C-t | *E. coli* BL21 | $1.80 \pm 0.59$ | 11 |
| | HNifU | 6xHis/N-t | *E. coli* BL21 | $1.89 \pm 0.04$ | 39 |
| | R-NifU<sub>s</sub> | Strep/C-t | *E. coli* BL21 | $5.54 \pm 0.45$ | 11 |
| | R-HNifU | 6xHis/N-t | *E. coli* BL21 | $6.80 \pm 0.93$ | 11 |
| | CA-HNifU | 6xHis/N-t | *E. coli* BL21 | $0.98 \pm 0.74$ | This work |
| | Apo-CA-HNifU | 6xHis/N-t | *E. coli* BL21 | $0.14 \pm 0.02$ | This work |
| | $\Delta C$ -HNifU | 6xHis/N-t | *E. coli* BL21 | $1.94 \pm 0.07$ | This work |
| | $\Delta N$ -HNifU | 6xHis/N-t | *E. coli* BL21 | $1.81 \pm 0.07$ | This work |
| NifS | sNifS | Strep/N-t | *E. coli* BL21 | - | 11 |
| NifH | Apo-NifH | Un-tagged | *A. vinelandii* | $0.34 \pm 0.11$ | 42,47 |
| | Holo-NifH | Un-tagged | *A. vinelandii* | $1.83 \pm 0.01$ | 42,47 |
| NifDK | Holo-NifDK | Un-tagged | *A. vinelandii* | $11.85 \pm 0.62^*$ | 42,47 |

Data are the average iron content per protein monomer ± SD calculated (n=3). S indicates Strep-tagged protein; H indicates 6xHis-tagged protein; R indicates proteins with *in vitro* reconstituted [Fe-S] clusters. \*The NifDK heterodimer was considered instead of a monomer of either protein.

**Figure 2.**
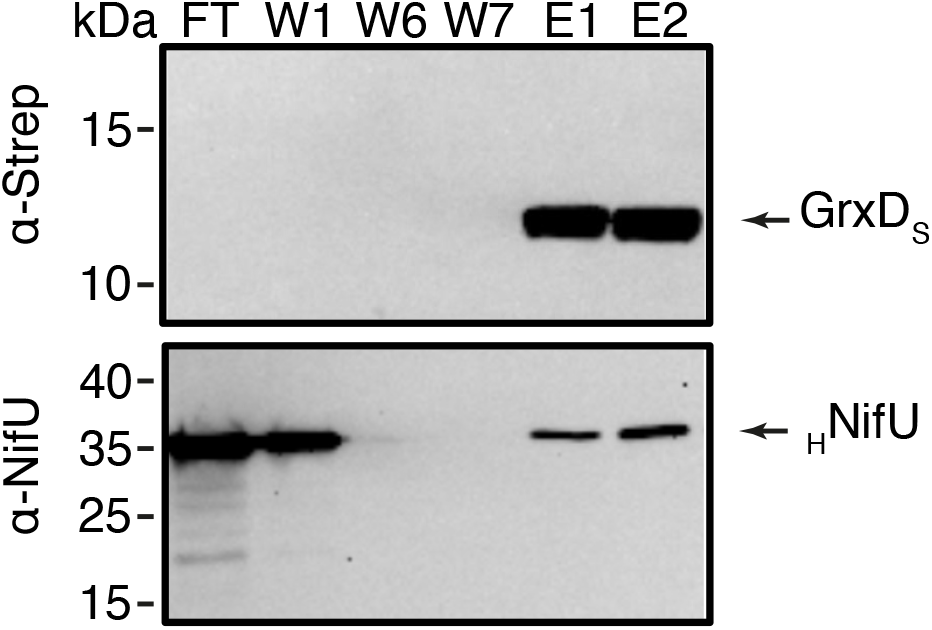
GrxD_S_ interacts with _H_NifU. Top panel shows the immunodetection with an anti-Strep antibody of Strep-tagged GrxD in flowthough (FT), washes (W1, W6, and W7), and elution (E1 and E2) fractions after being incubated with a histidine-tagged NifU and passed through a Strep-column. Bottom panel shows the immunoblot of the same fractions developed with an anti-NifU antibody. Images show a representative assay (n=3). Full length immunoblots are shown in Supplementary Figure S2.

To determine which domain is necessary for the interaction with GrxD, two NifU deletion mutants were produced: one without its N-domain (ΔN-NifU, containing amino acids 126-312, Table 2) and another without its C-domain (ΔC-NifU, containing amino acids 1-202, Table 2). Both proteins were fused to a N-terminal His-tag. Incubation of either protein with GrxD_S_, resulted in co-elution (Fig. 3, Fig. S5). As for wild type NifU, βN-_H_NifU, and βC-_H_NifU presence in the elution fraction was the result of their interaction with GrxD_S_ and not by any interaction with the resin on their own (Fig. S6). These interaction assays hint at NifU core domain as the main site interacting with GrxD, and likely acceptor of the [Fe_2_S_2_] cluster provided.

**Figure 3.**
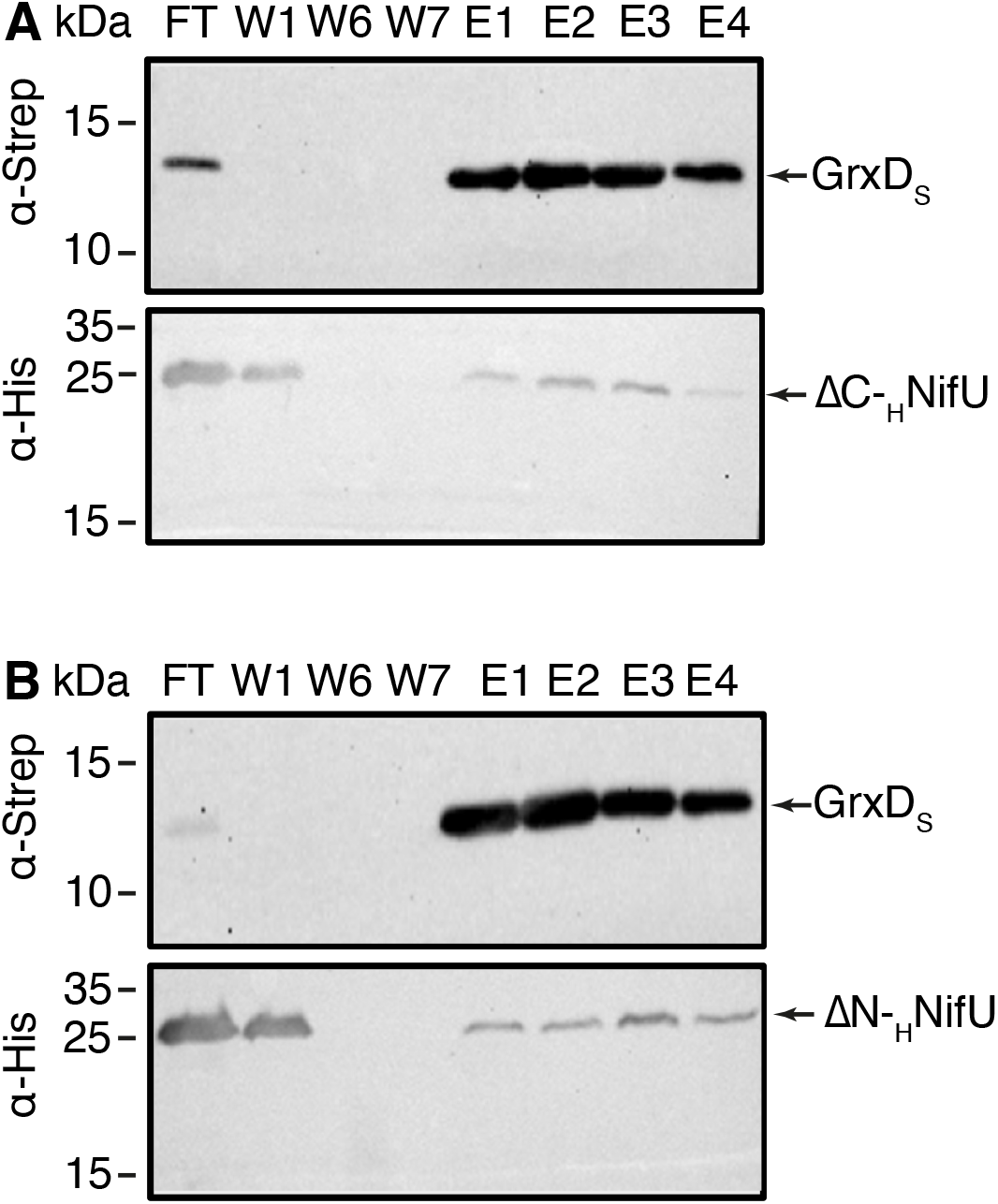
GrxD_S_ protein interacts with ΔN- and ΔC-_H_NifU mutants. A. Top panel shows the immunodetection with an anti-Strep antibody of Strep-tagged GrxD in flowthough (FT), washes (W1, W6, and W7), and elution (E1-E4) fractions after being incubated with a histidine-tagged ΔN-NifU and passed through a Strep-column. Bottom panel shows the immunoblot of the same fractions developed with an anti-His antibody. B. Top panel shows the immunodetection with an anti-Strep antibody of Strep-tagged GrxD in flowthough (FT), washes (W1, W6, and W7), and elution (E1-E4) fractions after being incubated with a histidine-tagged ΔC-NifU and passed through a Strep-column. Bottom panel shows the immunoblot of the same fractions developed with an anti-His antibody. Images show a representative assay (n=3). Full length immunoblots are shown in Supplementary Figure S5

### GrxD transfers a [Fe_2_S_2_] cluster to NifU

To rule out that GrxD could transfer [Fe_2_S_2_] groups to any of the transient sites in N- or C-domains, GrxD_S_ was incubated with _H_NifU, passed through a Strep-column, and iron and protein content determined in the flowthrough fraction. As shown in Fig. S7, no additional iron was observed in this fraction, indicating that no iron was transferred from GrxD_S_ to _H_NifU. Alternatively, iron could be transferred to the NifU core domain. This was explored by producing a N-terminal His-tagged NifU in which cysteines 35, 62, 106, 272, and 275 involved in the assembly of the [FeS] clusters at the N- and C-terminal domains were mutated to alanine (CA-_H_NifU; Table 2). The core [Fe_2_S_2_] group from this protein was fully removed by incubating the protein with urea and gently diluting it out to produce apo-CA-_H_NifU (Table 2). Incubating this protein with GrxD_S_, followed by separation in a Strep-column, resulted in 1.8 irons being transferred from GrxD to NifU (Fig. 4A, Fig. S8). This metal exchange was unidirectional, as reconstituted (R) NifU_S_ (Table 2) did not transfer any iron to apo-_H_GrxD (Fig. S9). Additionally, protein-protein interaction was required for metal transfer, since no iron was exchanged when the two proteins were separated by a dialysis membrane that only allowed for small molecule diffusion (Fig. 4B). This iron transfer was not due to dithiothreitol (DTT) being present in the buffer, as similar results were obtained when glutathione (GSH) was used instead (Fig. S10). UV-visible spectra of the resulting NifU were compatible with the transfer of the [Fe_2_S_2_] cluster (Fig. 4C). To confirm it, continuous-wave electron paramagnetic resonance (CW-EPR) spectroscopy was performed on the flow-through proteins (Figure 4D, Fig. S11), CA-_H_NifU showed a typical EPR signal for [Fe_2_-S_2_] (g = [2.02, 1.93, 1.89]) while GrxD_S_ did not exhibit an observable EPR signal, likely due to the presence of a diamagnetic [Fe_2_S_2_]^2+^ group, as previously reported (22, 23). Similarly, no cluster was observed in apo-CA-_H_NifU unless it was previously incubated with GrxD_S_, when a [Fe_2_-S_2_] group was detected.

**Figure 4.**
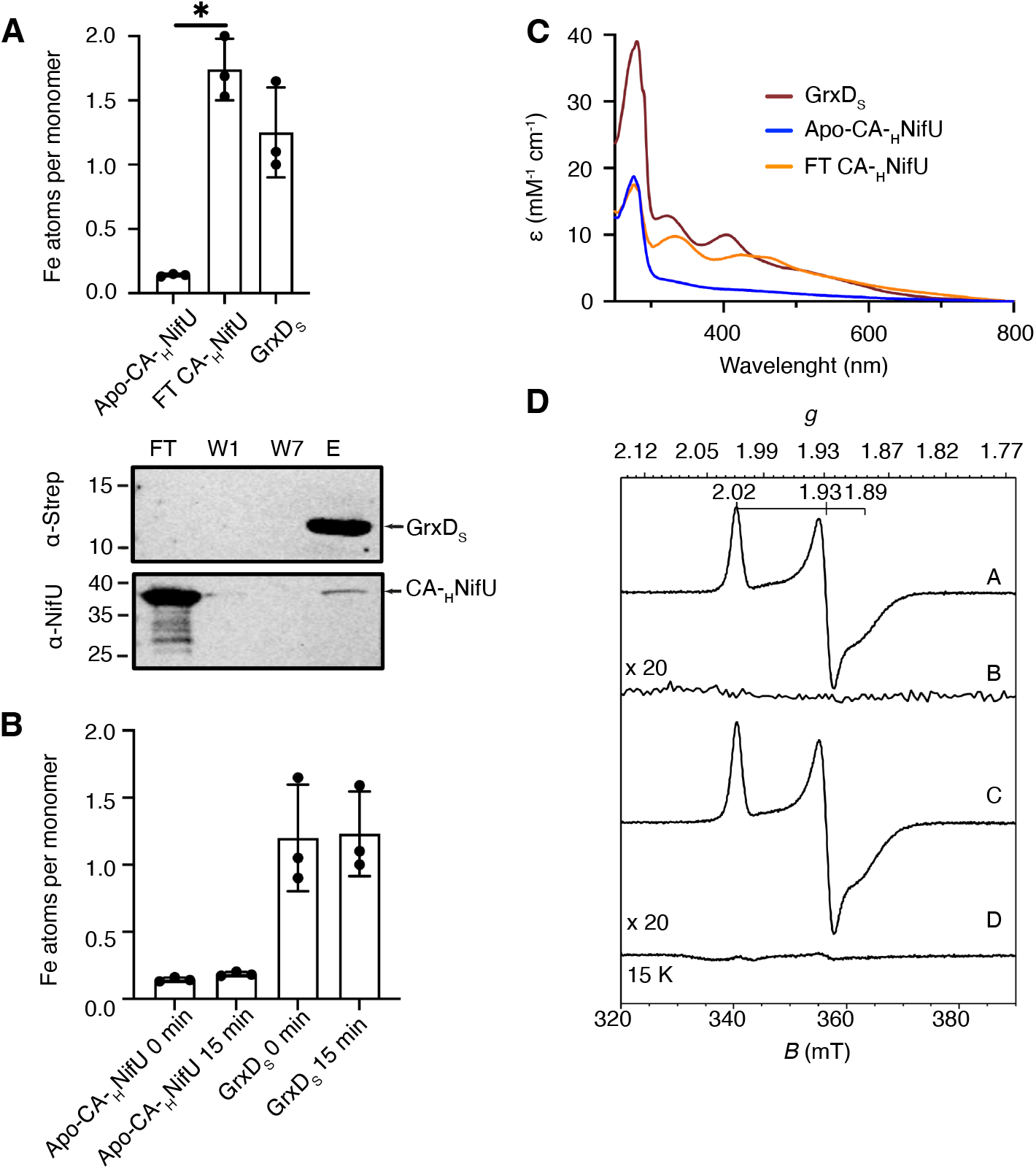
Apo-CA-_H_NifU receives a [Fe_2_S_2_] cluster from GrxD_S_. A. Top panel shows iron content of GrxD_S_ and apo-CA-_H_NifU prior to interaction, and in the flowthrough fraction (FT CA-_H_NifU) obtained from passing through a Strep-tactin-column a solution in which apo-CA-_H_NifU was incubated for 15 min with GrxD_S_. Bars represent the average ± SD (n=3). Bottom panel shows the immunoblots of flowthrough (FT), wash (W1-W7) and elution (E) fraction of the previous solution performed with anti-NifU and anti-Strep antibodies to confirm that not GrxD_S_ contamination was detected in the FT fraction. Images show a representative assay (n=3). Full length immunoblots are shown in Supplementary Figure S8. B. Iron content per monomer from proteins separated by a 3-kDa pore-size cutoff dialysis membrane before and after 15 min incubation. Bars represent the average ± SD (n=3). C. Molar extinction coefficients in UV-Visible spectra of the proteins in panel A. D. X-band cw-EPR spectra in the field range of 300-400 mT at 15 ºK under power-unsaturated conditions of the following samples: CA-_H_NifU (A) GrxD_S_ (B), flowthrough fraction after 15 min interaction of apo-CA-_H_NifU and AS-GrxD_S_ (C), and apo-CA-_H_NifU (D). A single S = ½ species with g = [2.02, 1.93, 1.89] is observed. The spin concentration of this species is 0.78 mM (A), 0 mM (B), 0.96 mM (C), and <5 µM (D). A second biological replicate is shown in Supplementary Figure S11B.

### GrxD delivers the [Fe_2_S_2_] catalytical cluster required for NifU activity

The previous results indicated that GrxD transfers a [Fe_2_S_2_] group to apo-CA-_H_NifU. However, they did not address whether CA-_H_NifU was properly refolded, kept the catalytic cluster, and re-gained the native NifU activity. *In vitro* NifU reconstitution and nitrogenase assays were used to validate the production of a functional NifU through GrxD-mediated [Fe_2_S_2_] transfer to apo-_H_NifU. NifU and NifS use free iron and cysteine to assemble the transient [Fe_4_S_4_] groups on NifU, resulting of approximately 6 Fe:NifU monomer (Fig. 5A) (9, 10). In contrast, when, apo-_H_NifU is combined with NifS, no additional irons were found on NifU, as it lacked the catalytic core cluster. However, should apo-_H_NifU be previously incubated with [Fe_2_S_2_]-containing GrxD_S_, repurified, and then combined with NifS, iron, and cysteine, the reconstitution of NifU is partially recovered. GrxD-mediated apo-NifU reactivation was also verified in *in vitro* acetylene reduction assays to determine nitrogenase activity. Maximal activity was achieved when using holo-NifH and holo-NifDK (Fig. 5B, Table 2). As expected, the activity was lost when holo-NifH was substituted by apo-NifH (Table 2), as electrons would not reach the catalytic site of NifDK. However, when R-_H_NifU (Table 2) was added to the mix, nitrogenase activity was restored to optimal levels, an activity that was not regained again if apo-_H_NifU was used instead. Validating the role of GrxD as provider of the NifU catalytic cluster, prior incubation of GrxD_S_ with apo-_H_NifU allowed the latter to synthesize and transfer [Fe_4_S_4_] clusters to apo-NifH leading to an active nitrogenase (Fig. 5B).

**Figure 5.**
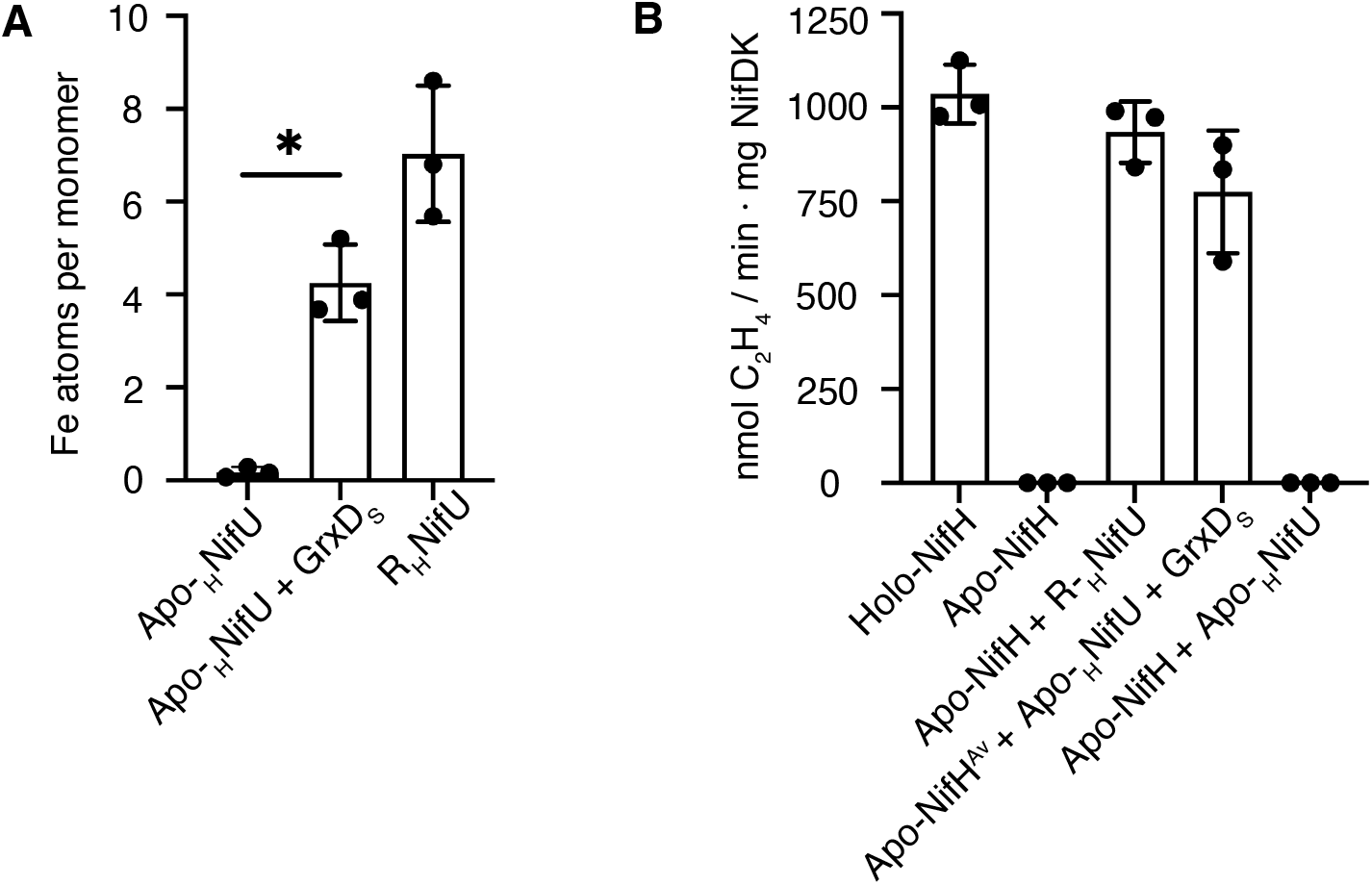
GrxD_S_ transfers the catalytic [Fe_2_S_2_] cluster to _H_NifU . A. *In vitro* synthesis of [FeS] clusters on _H_NifU by adding the reconstitution mix of apo-_H_NifU (1 mM L-cysteine, 1 mM DTT, 225 nM NifS, and 0.3 mM (NH_4_)_2_Fe(SO_4_)_2_). Three forms of _H_NifU were employed: as isolated _H_NifU (containing the catalytic [Fe_2_S_2_] in the core domain), apo-_H_NifU (lacking this cluster), and apo-_H_NifU that was previously incubated with GrxD_S_. B. Acetylene reduction assay using purified NifDK in combination with: holo-NifH, apo-NifH, apo-NifH combined with R-_H_NifU, apo-NifH combined with apo-_H_NifU previously incubated with GrxD_S_, and apo-NifH combined with apo-_H_NifU. Bars represent the average ± SD (n=3).

## Discussion

Glutaredoxins are a functionally diverse group of proteins participating in multiple cellular processes such as antioxidant defense, intracellular signaling and iron homeostasis (21, 24–26). In bacteria, they can be divided in two main groups: dithiol (Class I) and monothiol (Class II) Grxs (21). Class I can catalyze reversible thiol oxidation using glutathione as substrate (21). Monothiol Grxs do not have an enzymatic activity *per se* and act as [FeS] carriers (16, 17). These clusters are transferred to downstream [FeS] proteins such as IscA, ErpA, and enzymes involved in the membrane electron transport chain (27, 28). Our results extend the role of monothiol Grxs to delivering the catalytic [Fe_2_S_2_] cluster required by NifU to assemble the [Fe_4_S_4_] groups used to synthesize nitrogenase cofactors.

Among the three *A. vinelandii* Grxs, two of them belong to Class II: GrxD and Grx^nif^ (Avin51060). Both are up-regulated in the early stages of diazotrophy de-repression, when fixed nitrogen is depleted and *A. vinelandii* must start synthesizing nitrogenase. However, although Grx^nif^ is part of a *nif* cluster, no phenotype associated with nitrogen fixation has been reported to date (29). In contrast, GrxD presented a severe reduction of nitrogenase activity and overall iron content. This reduction led to lower growth rates under diazotrophic conditions, while no significant differences were observed under non-diazotrophic ones, even when iron was limiting in the medium. This contrasts with *E. coli* GrxD, that is required for adaptation to low-stress conditions (27). Therefore, some other mechanisms must be in place to accommodate the loss of GrxD function in *A. vinelandii* in non-diazotrophic conditions. The existence of functional alternatives to GrxD is also evidenced by the nitrogenase activity not being completely lost in *grxD* mutants.

While there are three [FeS] cluster binding sites in NifU, one in each of the three domains (8, 10), GrxD delivers exclusively a [Fe_2_S_2_] group to the core region. No iron transfer was observed from GrxD to core [Fe_2_S_2_]-containing NifU; only when this cluster was removed, iron could be moved from GrxD to apo-NifU or to apo-CA-NifU. Furthermore, this transfer was unidirectional, since apo-GrxD did not accept any iron from R-NifU. The polarity of metal cofactor donation seems to be a characteristic of many metal delivery systems as it prevents recovering essential, limiting, cofactors from metalloproteins. Such is the case of NifU and NifQ for [Fe_4_S_4_] (11), or CopZ and CopA for Cu^+^ (30), among others. Furthermore, iron delivery requires the docking of GrxD with NifU, as indicated by the lack of transfer when the two proteins are separated by a dialysis membrane that allows for diffusion of small molecules such as [Fe_2_S_2_] clusters, but not proteins. As a result, as in the case of other metallochaperones (31–33), metal delivery from GrxD to NifU is targeted, specific, and probably facilitated by the compatibility of the docking interfaces.

This biochemical data supports the phenotype observed in *grxD* mutants. Its reduced growth under diazotrophic conditions is the consequence of the lower nitrogenase activity of this strain. In turn, the lower performance of nitrogenase would stem from an altered NifU activity caused by GrxD deficiency. In this scenario, a large portion of NifU would lack the core [Fe_2_S_2_] group, hampering [Fe_4_S_4_] synthesis and transfer to Nif proteins as important as NifH, NifDK, NifB, and NifQ (8, 10–12). Consequently, nitrogenase would not receive its essential iron cofactors at the same rates, being less active overall. Considering that Nif proteins are a large part of *A. vinelandii* proteome during diazotrophy (18, 19, 29), many of them using iron cofactors (7), a reduction of [FeS] synthesis for these proteins could explain the lower iron concentration in *grxD* mutants.

Removing the [Fe_2_S_2_] cluster from NifU core domain required protein unfolding due to the high binding affinity of this region (34). However, addition of holo-GrxD allowed for proper refolding, as indicated by the recovery of the two irons lost during denaturing, as well as for the recovery of the [Fe_4_S_4_] synthesizing capability. This capability of functionally folding NifU could have a biotechnological role in current efforts to engineer nitrogen fixation capabilities in crop plants (35– 37). NifU has been shown to be an essential element for eukaryotes to produce NifH or NifB, as without it, the synthesis of the latter two proteins and their functionality *in vivo* are severely compromised (38, 39). However, NifU synthesis in plants seems to be limited by iron availability and, at most, NifU purified from plants contain one iron per monomer (40). Co-expression with GrxD could be an efficient way to optimize iron content in plant-produced NifU or to reduce the iron requirements in the nutrient solutions, in a similar way as it assists in NifU renaturing *in vitro*.

## Experimental procedures

### Escherichia coli *and* Azotobacter vinelandii *strains and plasmids*

*E. coli* strain BL21 (DE3) was used to express the proteins in this study. To produce GrxD_S_, *grxD* was amplified from *A. vinelandii* DNA using the primers 5NcoIGdxC-Strep and 3*Nde*IGdxC-Strep (Table S1), digested with *Nco*I and *Nde*I restriction enzymes and cloned in pET16bStrep to produce pERN1 (Table S2). _H_GrxD was generated by amplifying *GrxD* with primers 5*Nde*IGdxN-His and 3*Bam*HIGdxN-His digested with *Nde*I and *Bam*HI (Table S1) and ligated in pET16bHis, to assemble pERN2 (Table S2). ΔN-_H_NifUdomain, ΔC-_H_NifU, CA-_H_NifU encoding DNA were synthesized by GenScript (Piscataway, NJ, USA), flanked by *Nde*I and *Sex*AI restriction enzymes that were used to digest and ligate in pRHB612, originating plasmid pERN3, pERN4 and pERN5, respectively (Table S2).

To generate *A. vinelandii* strains, vector pDB2678 (Table S2) was produced by cloning a fragment comprising from -700 bp before *grxD* to 700bp after the stop codon. This fragment was amplifed by PCR using primers FW-*Not*I-700UpGrx5 and 700DownGrx5*Eco*RI-RV (Table S1), digested with *Not*I and *Eco*RI and ligated in pUC18. The in-frame *grxD* deletion mutant strain DJ3045 was produced by exchanging the wild type gene with an in-frame deletion of amino acids 2-50. This construction, pDB2679 (Table S2) was obtained by digesting and re-ligating the pDB2678 plasmid with *EcoRV*. Strain DC47 was obtained by exchanging this mutation with the wild type *grxD* clones in pDB2678. Transformation of *A. vinelandii* was carried out as described (41, 42). To select for the transformants, the strains were co-transformed with pDB303 (to select with 10 μg/mL Rifampicin for DJ3045) and pDB1416 (to select with 0.05 μg/mL Gentamycin for DC47) provided by Dr. Dennis Dean (Virginia Tech, Blacksburg, USA).

#### Protein purification

Protein synthesis in *E. coli* was induced by adding 1 mM isopropyl β-D-thiogalactoside (IPTG) to cells growing in LB media supplemented with 100 mg/ml ampicillin at OD_600_ ≈ 0.6. After 6 h of induction at 30 ºC, cells were collected by centrifugation at 4000 *g* for 7 min and stored at -80 ºC. Cells producing _H_NifU_S_ or _H_GrxD_S_ were further supplemented with 0.2 mM ferric ammonium citrate and 2 mM L-cysteine. In the case of cells synthesizing _S_NifS, the medium was supplemented with 2 mM L-cysteine. Strep-tagged proteins were purified by Strep-Tactin XT affinity chromatography. Approximately 15 to 20 g of cells were resuspended for 30 min in 60 ml of Buffer W (100 mM Tris-HCl pH 8.0, 150 mM NaCl) containing 1 mM phenyl-methylsulfonyl fluoride (PMSF) and 3 mg of DNAse I. Cells were lysed in a French Press cell at 1500 lb per square inch. The cell-free extract (CFE) was obtained after removing cell debris by centrifugation at 63,000 *g* for 1 h at 4 º C. CFE was loaded onto a 3 ml Gravity flow Streptactin-XT high-capacity column (IBA, Göttingen, Germany), previously equilibrated with buffer W. The column was then washed 5 times with two column-volumes (CV) of buffer W per wash. The bound proteins were eluted with 5 CV of 50 mM biotin in buffer W.

His-tagged proteins were purified by Ni-NTA affinity chromatography. Approximately 20 to 25 g of *E. coli* BL21(DE3) cells were resuspended for 30 min in 100 ml of buffer A (100 mM Tris-HCl pH 8.0, 250 mM NaCl, and 20 mM imidazole) supplemented with 1 mM PMSF and 3 mg DNAse I. Cells were lysed and CFE was obtained as described above. CFE was loaded onto a 5 ml Ni-NTA Agarose column (Qiagen, Hilden, Germany) previously equilibrated with buffer A. The column was washed with 10 CV of 20 mM imidazole in buffer A with and with 10 CV of 40 mM imidazole in buffer A. In the case of CA-NifU, a third wash was performed with 90 mM of imidazole in buffer A. Elutions were performed with 300 mM imidazole in buffer A.

NifH and NifDK were produced and purified from *A. vinelandii* DJ (NifDK) and DJ33 (NifH) as described by Shah VK et al. (42)

Elution fractions were concentrated with centrifugal membrane devices (Amicon Ultra, Millipore, Burlington, MA, USA). The proteins were desalted using a PD-10 column previously equilibrated in Buffer W. All purifications were carried out under anaerobic conditions (<5.0 ppm O_2_) inside a glovebox (COY Laboratories, Grass Lake, MI, USA) using buffers previously made anaerobic by sparging with N_2_ overnight. Purification fractions were analyzed by electrophoresis. Purified proteins were frozen and stored in liquid N_2_. Protein and iron content was determined using BCA (Pierce, IL, USA) and bipyridyl method (43), respectively.

### In vitro *[FeS] cluster reconstitution*

[FeS] cluster reconstitution on NifU was carried out as described (11). Briefly, 20 μM of NifU dimer was prepared in 100 mM MOPS (pH 7.5) buffer containing 8 mM 1,4-dithiothreitol (DTT) and incubated at room temperature for 30 min. To this solution, 1 mM L-cysteine, 1 mM DTT, 225 nM NifS, and 0.3 mM (NH_4_)_2_Fe(SO_4_)_2_ were added. Iron additions were divided in three steps of 15 min each until reaching the final concentration of 0.3 mM. This solution was maintained in ice for 3 h. To remove unbound iron, R-NifU was desalted by dilution and concentrated with a 10-kDa cutoff pore size centrifugal membrane device (Millipore, Burlington, MA, USA) in 5 mM DTT in 100 mM MOPS (pH 7.4). R-NifU protein was stored in liquid N_2_ until use. This work was carried out under anaerobic conditions (<5.0 ppm O_2_) inside a glovebox using buffers previously made anaerobic by sparging with N_2_ overnight. Protein and iron content was determined using Bradford (Pierce) and bipyridyl method (43), respectively.

#### [FeS] cluster removal

To remove all the [FeS] clusters from CA-_H_NifU and _H_NifU, 100 μM of these proteins were incubated at room temperature for 4 h in 100 mM Tris (pH 8.0), 150 mM NaCl, 8 M Urea, 100 mM EDTA, and 5 mM DTT. The protein was desalted in decreasing Urea solutions (6 M, 4 M, 2 M and 0.5 M) in Buffer W using a 10-kDa cutoff pore size centrifugal membrane device. Finally, it was passed through a PD-10 previously equilibrated in Buffer EPR (100 mM Tris-HCl pH 8.0, 350 mM NaCl, 10% glycerol). The proteins were stored in liquid N_2_ until use.

The clusters from _H_GrxD were eliminated by incubating 50 μM of this protein in 50 μM of EDTA in Buffer W for 1 h at room temperature. The solution was concentrated using a 3-kDa cutoff pore size centrifugal membrane devices (Millipore). Finally, it was passed through a PD-10 for desalting in Buffer W. Apo-_H_GrxD was stored in liquid N_2_ until use.

Apo-NifH was prepared by incubated 30 min at room temperature in 22 mM Tris-HCl pH 7.4, 2 mM DTH, 2.5 mM ATP, 2.5 mM MgCl_2_ and 40 mM 2,2’ Bipyridine. The protein mix was desalting using a PD-10 column previously equilibrated in 100 mM Tris-HCl pH 8.0, 200 mM NaCl, 10% glycerol and 2 mM DTH, the elution was concentrated using a 10 kDa cutoff pore size centrifugal membrane devices (Millipore). Apo-NifH was stored in liquid N_2_ until use.

#### Co-purification assays

All interaction assays between GrxD and NifU were carried out for 5 min using 20 μM of each protein in a final volume of 200 μl of Buffer W inside a glovebox under anaerobic conditions. The proteins were separated using a 200 μl of Gravity flow Streptactin-XT high-capacity (IBA) column. The column was washed seven times with 3 CV of Buffer W. The elution was carried out in 4 steps using 2 CV of 50 mM biotin in Buffer W.

#### [FeS] cluster transfer assays

To determine iron transfer between apo-CA-_H_NifU or apo-_H_NifU and GrxD_S_, a combination of 50 μM of the NifU partner and 250 μM of the GrxD pair were incubated in a glovebox for 15 min at room temperature in EPR buffer. Proteins were separated by passing them through a column of 5 ml Strep-tactin XT 4Flow High-capacity column (IBA) previously equilibrated in buffer W. The column was washed with seven CVs of buffer W, and the proteins were eluted with 50 mM biotin in Buffer W. To test the diffusion of free iron from reconstituted proteins to apo-protein, the two proteins were incubated for 15 min inside an anaerobic glovebox, separated by a dialysis membrane. Samples from both sides of the membrane were collected to determine the protein and iron concentration at time 0 and after 15 min incubation. To prepare EPR samples, the flowthrough fraction was concentrated using 30 kDa cutoff pore size centrifugal membrane devices (Millipore) and stored in liquid N_2_

Protein content in all selected fractions was analyzed by SDS-PAGE using 15% acrylamide/bisacrylamide (37.5:1) gels and visualized by Coomassie Brilliant Blue staining (44). For immunoblot analysis, proteins were transferred to nitrocellulose membranes for 30 min at 20 V using a Transfer-Blot Semi Dry system (Bio-Rad, Hercules, CA, USA). Immunoblot analyses were carried out with primary mouse antibodies raised against Strep tag (1:2,500 dilution Strep-MAB) (IBA) or against (His)_6_ tag (1:2,000 dilution α-His primary monoclonal antibody) (Sigma-Aldrich, St Louis, MO, USA), and a secondary anti-mouse antibody conjugated to peroxidase (1:10,000 dilution, Goat anti-Mouse IgG) (Agrisera, Vännäs, Sweden). In the case of NifU, a primary specific anti-NifU rabbit antibody was used at 1:2,500 dilution (45) in combination with a secondary horseradish peroxidase-conjugated anti-rabbit antibody (IBA) diluted 1:15,000. Chemiluminescent detection was carried out according to Pierce ECL Western Blotting Substrate kit’s instructions (ThermoFisher Scientific, Waltham, MA, USA) and developed in an iBright FL1000 Imaging System (ThermoFisher Scientific). Protein content was determined using BCA (Pierce, IL, USA) and iron content with Atomic Absorption Spectroscopy.

#### Iron determinations with atomic absorption spectroscopy

Protein samples were mineralized in 37.5 % analytic grade nitric acid for 10 min at 80 ºC. Samples were then diluted to a total concentration of 2 % nitric acid with ultrapure water. To determine iron content in cellular samples, 1 ml of the cultures was pelleted and digested in 50 μl 6 N HCl overnight. Samples were diluted 10-fold to reduce acid concentrations. Iron concentrations were determined in an Atomic Absorption Spectrometer ContrAA 800G (AnalytikJena, Jena, Germany) using commercially available analytic grade metal standards (Inorganic Ventures, Christianburg, VA, USA).

#### Ultraviolet-visible spectroscopy

UV-visible absorption spectra were collected under anaerobic conditions (<0.1 ppm O_2_) inside a glovebox (MBraun, München, Germany) in septum-sealed cuvettes to avoid the O_2_ contamination during the measurements in the Shimadzu UV-2600 spectrophotometer. Absorption (225 nm to 800 nm) was recorded.

#### Electron paramagnetic resonance (EPR) spectroscopy

Protein samples were prepared in EPR buffer supplemented with 1 mM dithionite (DTH). X-band (9.64 GHz) cw-EPR spectra were recorded on a Bruker Elexsys spectrometer equipped with an Oxford ESR 910 cryostat and a Bruker cavity. The microwave frequency was calibrated with a frequency counter and the magnetic field with an NMR gauss meter. The temperature was calibrated with a carbon-glass resistor temperature probe (CGR-1-1000; Lake-Shore Cryotronics) located into an EPR tube. For all EPR spectra, a modulation frequency and amplitude of 100 kHz and 1 mT were used. The first-derivative spectra were recorded at 1024 points with an integration time of 150 milliseconds. EPR spectral simulations were performed using the simulation software SpinCount (46). The spin was quantified by relative to a 1.2 mM Cu(II)ethylenediaminetetraacetic solution with 10% glycerol (v/v). Two EPR samples, independently prepared from two different GrxD_s_ and CA-_H_NifU interaction assays, were measured.

#### Nitrogenase activity assays

Prior to culture, all flasks and vials were acid-washed overnight in 0.2 M HCl to remove iron contaminations, followed by thoroughly rinsing in deionized water. 250 ml cultures of each *A. vinelandii* strain were grown overnight at 30ºC 200 rpm in 250 ml of Burk medium (41). On the following day, these cultures were centrifuged at 1400 *g* 10 minutes and resuspended with a OD_600_ 0.3 in 250 ml of Burk medium for 2 hours at 30ºC with 200 rpm agitation. To de-repress nitrogenase, the cells were washed with Burk Medium without any nitrogen source by centrifugation and resuspended again in 50 ml of the same medium and grown for 4 h at 30 ºC with 200 rpm agitation. Nitrogenase activity was determined with the acetylene reduction assay (47). One milliliter of culture was incubated with 0.5 ml of acetylene for 15 minutes at 30 ºC in 9 ml vials closed with rubber caps. Reaction was stopped using 0.1 ml 8 M NaOH. Two samples of 50 μl were analyzed in gas chromatographer GC-8A (Shimadzu, Kyoto, Japan) using a Porapak N 80/100 column. The ethylene production rates were normalized to the OD_600_ of the cultures after 4 hours of diazotrophic growth.

*In vitro* reconstitution of apo-NifH by NifU was determined as described (40, 48). Reactions were prepared inside anaerobic chambers. 4 μM of apo-NifH was incubated with 0.1 μM of NifDK, 40 μM NifU and ATP-regenerating mixture (1.23 mM ATP, 18 mM phosphocreatine, 2.2 mM MgCl_2_, 3 mM DTH, 5 mM DTT, 46 μg/ml of creatine phosphokinase, 100 mM MOPS pH 7.4) in a final volume of 600 μl inside 9 ml serum vials under argon atmosphere containing 500 μl acetylene (1 atm). The reaction was carried out at 30 °C in a shaking water bath for 15 min. Reactions were stopped by adding 100 μl of 8 M NaOH. Ethylene formed was measured in 50 μl gas phase samples as previously indicated.

#### Statistical methods

GraphPad Prism software was used for statistical analysis. The data were compared using unpaired t test with Welch’s correction (*p* <0.05)

## Data Availability

The authors declare that the data supporting the findings of this study are available within the article, its supplementary information and data, and upon request.

## Supporting Information

This article contains supporting information.

## Acknowledgements

The authors would like to acknowledge Dr. Emma Barahona and Mr. Álvaro Salinero (CBGP, UPM-INIA/CSIC) for their help in *A. vinelandii* transformations; Dr. Dennis Dean (Virginia Tech, USA) for providing vectors; and Dr. Jin Xiong (Carnegie Mellon University, USA) for his help in the EPR measurements.

## Author contribution

JAC-G, LMR, and MG-G writing, review, editing, and formal analyses; LMR and MG-G supervision, conceptualization, and funding acquisition; JAC-G, ER-N, AMA, DR, AP-G, YG, and CE-E, investigation.

## Funding and additional information

This work was by grant PID2021-12460OB-100 from the Ministerio de Ciencia, Innovación/Agencia Estatal de Investigación/10.13039/50110001103 and “ERDF A way of making Europe to MG-G. This work was supported by the Bill and Melinda Gates Foundation grant INV-005889 and the Gates Foundation grant INV-067006 to LMR. Under the grant conditions of the Foundation, a Creative Commons Attribution 4.0 Generic License has already been assigned to the author accepted manuscript version that might arise from this submission. YG acknowledges the funding supporting from Department of Energy (DOE-DE-SC0026054). JAC-G and ER-N were supported by Formación de Personal de Investigación fellowships PRE2022-101253 and PRE2018-084895, respectively. The latter was part of the Severo Ochoa Programme for Centres of Excellence in R&D from Agencia Estatal de Investigación (grants SEV-2016-0672 and CEX2020-000999S) received by Centro de Biotecnología y Genómica de Plantas (UPM-INIA/CSIC). AMA was funded by a María Zambrano contract. AP-G is a recipient of a Ramón y Cajal grant (RYC2021-031246-I) funded by MCIN/AEI/10.13039/501100011033 and by the European Union NextGenerationEU/PRTR.

## References

1. Hoffman, B. M., Lukoyanov, D., Yang, Z.-Y., Dean, D. R., and Seefeldt, L. C. (2014) Mechanism of nitrogen fixation by nitrogenase: The next stage. Chem. Rev. 114, 4041–4062

2. Bulen, W. A., and LeComte, J. R. (1966) The nitrogenase system from Azotobacter: two-enzyme requirement for N2 reduction, ATP-dependent H2 evolution, and ATP hydrolysis. Proc. Natl. Acad. Sci. U S A. 56, 979–986

3. Einsle, O., Tezcan, F. A., Andrade, S. L. A., Schmid, B., Yoshida, M., Howard, J. B., and Rees, D. C. (2002) Nitrogenase MoFe-Protein at 1.16 Å resolution: A central ligand in the FeMo-cofactor. Science 297, 1696 –1700

4. Spatzal, T., Aksoyoglu, M., Zhang, L., Andrade, S. L. A., Schleicher, E., Weber, S., Rees, D. C., and Einsle, O. (2011) Evidence for interstitial carbon in nitrogenase FeMo Cofactor. Science 334, 940 –940

5. Seefeldt, L. C., Peters, J. W., Beratan, D. N., Bothner, B., Minteer, S. D., Raugei, S., and Hoffman, B. M. (2018) Control of electron transfer in nitrogenase. Curr. Opin. Chem. Biol. 47, 54–59

6. Rutledge, H. L., and Tezcan, F. A. (2020) Electron transfer in nitrogenase. Chem. Rev. 120, 5158–5193

7. Burén, S., Jiménez-Vicente, E., Echavarri-Erasun, C., and Rubio, L. M. (2020) Biosynthesis of Nitrogenase Cofactors. Chem. Rev. 120, 4921–4968

8. Dos Santos, P. C., Smith, A. D., Frazzon, J., Cash, V. L., Johnson, M. K., and Dean, D. R. (2004) Iron-Sulfur Cluster Assembly: NifU-directed activation of the nitrogenase Fe protein. J. Biol. Chem. 279, 19705–19711

9. Yuvaniyama, P., Agar, J. N., Cash, V. L., Johnson, M. K., and Dean, D. R. (2000) NifS-directed assembly of a transient [2Fe-2S] cluster within the NifU protein. Proc. Natl. Acad. Sci. USA. 97, 599–604

10. Smith, A. D., Jameson, G. N. L., Dos Santos, P. C., Agar, J. N., Naik, S., Krebs, C., Frazzon, J., Dean, D. R., Huynh, B. H., and Johnson, M. K. (2005) NifS-Mediated Assembly of [4Fe−4S] clusters in the N- and C-terminal domains of the NifU scaffold protein. Biochemistry. 44, 12955–12969

11. Barahona, E., Collantes-Garcia, J., Rosa-Núñez, E., Xiong, J., Jiang, X., Jiménez-Vicente, E., Echávarri-Erasun, C., Guo, Y., Rubio, L., and González-Guerrero, M. (2024) Azotobacter vinelandii scaffold protein NifU transfers iron to NifQ as part of the iron-molybdenum cofactor biosynthesis pathway for nitrogenase. J. Biol. Chem. 300, 107900

12. Zhao, D., Curatti, L., and Rubio, L. M. (2007) Evidence for nifU and nifS Participation in the Biosynthesis of the Iron-Molybdenum cofactor of nitrogenase. J. Biol. Chem. 282, 37016–37025

13. Schmitz, R. A., Klopprogge, K., and Grabbe, R. (2002) Regulation of nitrogen fixation in Klebsiella pneumoniae and Azotobacter vinelandii: NifL, transducing two environmental signals to the nif transcriptional activator NifA. J. Mo.l Microbiol. Biotechnol. 4, 235–42

14. Zheng, L., Cash, V. L., Flint, D. H., and Dean, D. R. (1998) Assembly of Iron-Sulfur clusters. Identification of an iscSUA-hscBA-fdx gene cluster from Azotobacter vinelandii. J. Biol. Chem. 273, 13264–13272

15. Shakamuri, P., Zhang, B., and Johnson, M. K. (2012) Monothiol glutaredoxins function in storing and transporting [Fe2 S2] clusters assembled on IscU scaffold proteins. J. Am. Chem. Soc. 134, 15213–15216

16. Li, H., Mapolelo, D. T., Dingra, N. N., Naik, S. G., Lees, N. S., Hoffman, B. M., Riggs-Gelasco, P. J., Huynh, B. H., Johnson, M. K., and Outten, C. E. (2009) The yeast iron regulatory proteins Grx3/4 and Fra2 form heterodimeric complexes containing a [2Fe-2S] cluster with cysteinyl and histidyl ligation. Biochemistry 48, 9569–9581

17. Roret, T., Tsan, P., Couturier, J., Zhang, B., Johnson, M. K., Rouhier, N., and Didierjean, C. (2014) Structural and spectroscopic insights into BolA-Glutaredoxin complexes. J. Biol. Chem. 289, 24588–24598

18. Rosa-Núñez, E., Echavarri-Erasun, C., Armas, A. M., Escudero, V., Poza-Carrión, C., Rubio, L. M., and González-Guerrero, M. (2023) Iron homeostasis in Azotobacter vinelandii. Biology 12, 1423

19. Poza-Carrión, C., Jiménez-Vicente, E., Navarro-Rodríguez, M., Echavarri-Erasun, C., and Rubio, L. M. (2014) Kinetics of nif gene expression in a nitrogen-fixing bacterium. J. Bacteriol. 196, 595–603

20. Fu, W., Jack, R. F., Morgan, T. V, Dean, D. R., and Johnson, M. K. (1994) nifU gene product from Azotobacter vinelandii is a homodimer that contains two identical [2Fe-2S] clusters. Biochemistry 33, 13455–13463

21. Talib, E. A., and Outten, C. E. (2021) Iron-sulfur cluster biogenesis, trafficking, and signaling: Roles for CGFS glutaredoxins and BolA proteins. Biochim. Biophys. Acta Mol. Cell. Res. 1868, 118847

22. Lillig, C. H., Berndt, C., Vergnolle, O., Lönn, M. E., Hudemann, C., Bill, E., and Holmgren, A. (2005) Characterization of human glutaredoxin 2 as iron-sulfur protein: a possible role as redox sensor. Proc. Natl. Acad. Sci. USA. 102, 8168–73

23. Rouhier, N., Unno, H., Bandyopadhyay, S., Masip, L., Kim, S.-K., Hirasawa, M., Gualberto, J. M., Lattard, V., Kusunoki, M., Knaff, D. B., Georgiou, G., Hase, T., Johnson, M. K., and Jacquot, J.-P. (2007) Functional, structural, and spectroscopic characterization of a glutathione-ligated [2Fe–2S] cluster in poplar glutaredoxin C1. Proc. Natl. Acad. Sci. USA. 104, 7379–7384

24. Herrero, E., and de la Torre-Ruiz, M.A. (2007) Monothiol glutaredoxins: a common domain for multiple functions. Cell. Mol. Life Sci. 64, 1518–1530

25. Hati, D., Brault, A., Gupta, M., Fletcher, K., Jacques, J.-F., Labbé, S., and Outten, C. E. (2023) Iron homeostasis proteins Grx4 and Fra2 control activity of the Schizosaccharomyces pombe iron repressor Fep1 by facilitating [2Fe-2S] cluster removal. J. Biol. Chem. 299, 105419

26. Ogata, F. T., Branco, V., Vale, F. F., and Coppo, L. (2021) Glutaredoxin: Discovery, redox defense and much more. Redox Biol. 43, 101975

27. Fisher, C. E., Bak, D. W., Miller, K. E., Washington-Hughes, C. L., Dickfoss, A. M., Weerapana, E., Py, B., and Outten, F. W. (2024) Escherichia coli monothiol glutaredoxin GrxD replenishes Fe-S clusters to the essential ErpA A-type carrier under low iron stress. J. Biol. Chem. 300, 107506

28. Azam, T., Przybyla-Toscano, J., Vignols, F., Couturier, J., Rouhier, N., and Johnson, M. K. (2020) The Arabidopsis mitochondrial glutaredoxin GRXS15 provides [2Fe-2S] clusters for ISCA-mediated [4Fe-4S] cluster maturation. Int. J. Mol. Sci. 21, 9237

29. Martin del Campo, J.S., Rigsbee, J., Bueno Batista, M., Mus, F., Rubio, L. M., Einsle, O., Peters, J. W., Dixon, R., Dean, D. R., and Dos Santos, P. C. (2022) Overview of physiological, biochemical, and regulatory aspects of nitrogen fixation in Azotobacter vinelandii. Crit. Rev. Biochem. Mol. Biol. 57, 492–538

30. González-Guerrero, M., and Argüello, J. M. (2008) Mechanism of Cu^+^-transporting ATPases: Soluble Cu^+^-chaperones directly transfer Cu^+^ to transmembrane transport sites. Proc. Natl. Acad. Sci. U S A. 105, 5992–5997

31. Navarro-Gómez, C., León-Mediavilla, J., Küpper, H., Rodríguez-Simón, M., Paganelli-López, A., Wen, J., Burén, S., Mysore, K. S., Bokhari, S. N. H., Imperial, J., Escudero, V., and González-Guerrero, M. (2024) Nodule-specific Cu^+^-chaperone NCC1 is required for symbiotic nitrogen fixation in Medicago truncatula root nodules. New Phytol. 241, 793–810

32. Patel, S. J., Frey, A. G., Palenchar, D. J., Achar, S., Bullough, K. Z., Vashisht, A., Wohlschlegel, J. A., and Philpott, C. C. (2019) A PCBP1–BolA2 chaperone complex delivers iron for cytosolic [2Fe–2S] cluster assembly. Nat. Chem. Biol. 15, 872–881

33. Weiss, A., Murdoch, C. C., Edmonds, K. A., Jordan, M. R., Monteith, A. J., Perera, Y. R., Rodríguez Nassif, A.M., Petoletti, A. M., Beavers, W. N., Munneke, M. J., Drury, S. L., Krystofiak, E. S., Thalluri, K., Wu, H., Kruse, A. R. S., DiMarchi, R. D., Caprioli, R. M., Spraggins, J. M., Chazin, W. J., Giedroc, D. P., and Skaar, E. P. (2022) Zn-regulated GTPase metalloprotein activator 1 modulates vertebrate zinc homeostasis. Cell. 185, 2148-2163.e27

34. Agar, J. N., Yuvaniyama, P., Jack, R. F., Cash, V. L., Smith, A. D., Dean, D. R., and Johnson, M. K. (2000) Modular organization and identification of a mononuclear iron-binding site within the NifU protein. J. Biol. Inorg. Chem. 5, 167–177

35. Burén, S., and Rubio, L. M. (2018) State of the art in eukaryotic nitrogenase engineering. FEMS Microbiol. Lett. 365, fnx274–fnx274

36. Mus, F., Crook, M. B., Garcia, K., Garcia Costas, A., Geddes, B. A., Kouri, E. D., Paramasivan, P., Ryu, M. H., Oldroyd, G. E., Poole, P. S., Udvardi, M. K., Voigt, C. A., Ane, J. M., and Peters, J. W. (2016) Symbiotic nitrogen fixation and challenges to extending it to non-legumes. Appl. Environ. Microbiol. 82, 3698–3710

37. Zhao, Z., Fernie, A. R., and Zhang, Y. (2025) Engineering nitrogen and carbon fixation for next-generation plants. Curr. Opin. Plant Biol. 85, 102699

38. Eseverri, A., López-Torrejón, G., Jiang, X., Buren, S., Rubio, L., and Caro, E. (2020) Use of synthetic biology tools to optimize the production of active nitrogenase Fe protein in chloroplasts of tobacco leaf cells. Plant Biotechnol. J. 18, 1882–1896

39. Buren, S., Pratt, K., Jiang, X., Guo, Y., Jiménez-Vicente, E., Echavarri-Erasun, C., Dean, D., Saaem, I., Gordon, D., Voight CA, and Rubio, L. (2019) Biosynthesis of the nitrogenase active-site cofactor precursor NifB-co in Saccharomyces cerevisiae. Proc. Natl. Acad. Sci. U S A. 116, 25078–25086

40. Jiang, X., Payá-Tormo, L., Coroian, D., García-Rubio, I., Castellanos-Rueda, R., Eseverri, Á., López-Torrejón, G., Burén, S., and Rubio, L. M. (2021) Exploiting genetic diversity and gene synthesis to identify superior nitrogenase NifH protein variants to engineer N2-fixation in plants. Commun Biol. 4, 4

41. Dos Santos, P. C. (2019) Genomic manipulations of the diazotroph Azotobacter vinelandii. Methods Mol. Biol. 1876, 91–109

42. Shah, V. K., and Brill, W. J. (1973) Simple method of purification to homogeneity of nitrogenase components from Azotobacter vinelandii. Biochim Biophys Acta. 305, 445–54

43. Moss, M. L., and Mellon, M. G. (1942) Colorimetric determination of iron with 2,2’-Bipyridyl and with 2,2’,2’-Terpyridyl. Ind. Eng. Chem. Anal. Ed. 14, 862–865

44. Laemmli, U. K. (1970) Cleavage of structural proteins during the assembly of the head of bacteriophage T4. Nature 227, 680–685

45. Lopez-Torrejon, G., Jimenez-Vicente, E., Buesa, J. M., Hernandez, J. A., Verma, H. K., and Rubio, L. M. (2016) Expression of a functional oxygen-labile nitrogenase component in the mitochondrial matrix of aerobically grown yeast. Nat. Commun. 7, 11426

46. Petasis, D. T., and Hendrich, M. P. (2015) Quantitative interpretation of multifrequency multimode EPR spectra of metal containing proteins, enzymes, and biomimetic complexes. Methods Enzymol. 563, 171–208

47. Robinson, A. C., Burgess, B. K., and Dean, D. R. (1986) Activity, reconstitution, and accumulation of nitrogenase components in Azotobacter vinelandii mutant strains containing defined deletions within the nitrogenase structural gene cluster. J. Bacteriol. 166, 180–186

48. Shah, V. K., Imperial, J., Ugalde, R. A., Ludden, P. W., and Brill, W. J. (1986) In vitro synthesis of the iron-molybdenum cofactor of nitrogenase. Proc. Natl. Acad. Sci. USA. 83, 1636–1640

49. Setubal, J. C., dos Santos, P., Goldman B.S., Erstevag H., Espin G., et al. (2009). Genome sequence of Azotobacter vinelandii, an obligate aerobe specialized to support diverse anaerobic metabolic processes. J. Bacteriol. 191, 4534–4545

