## Supplemental for "*Azotobacter vinelandii* glutaredoxin D delivers the core [Fe_2_S_2_] cluster to nitrogenase cofactor scaffold protein NifU": Supplemental Figure Legends.pdf

**Figure S1. Mutation in *grxD* does not affect non-diazotrophic growth under low iron conditions.** Growth under non-diazotrophic, low iron conditions of wild type *A. vinelandii* strain (DJ), *grxD* in-frame mutant (DJ3045), and DJ3045 transformed with a wild-type copy of *grxD* (DC47). Bars represent the average ± SD (n=3).

**Figure S2. Full length immunoblots shown in Figure 2.**

**Figure S3. $_{H}$NifU does not bind to the Strep-column.** Immunodetection with an anti-NifU antibody of His-tagged NifU in flowthough (FT), washes (W1, W6, and W7), and elution (E1 and E2) fractions after passing through a Strep-column. Images show a representative assay (n=3).

**Figure S4. GrxD and NifU proteins interacts independently of their tags.** A. Immunodetection with an anti-His antibody of His-tagged GrxD in flowthough (FT), washes (W1, W6, and W7), and elution (E1 and E2) fractions after incubation with Strep-tagged NifU and passed through a Strep-column. B. Same fractions as above but visualized with an anti-Strep antibody. C. Immunodetection with an anti-His antibody of His-tagged GrxD in flowthough (FT), washes (W1, W6, and W7), and elution (E1 and E2) fractions after passing through a Strep-column. Images show a representative assay (n=3).

**Figure S5. Full length immunoblots shown in Figure 3.**

**Figure S6. ΔN and ΔC $_{H}$NifU donot bind to a Strep-column.** A. Immunodetection with an anti-His antibody of His-tagged ΔN-NifU in flowthough (FT), washes (W1, W6, and W7), and elution (E1-E4) fractions after passing through a Strep-column. B. Immunodetection with an anti-His antibody of His-tagged ΔC-NifU in flowthough (FT), washes (W1, W6, and W7), and elution (E1-E4) fractions after passing through a Strep-column. Images show a representative assay (n=3).

**Figure S7. GrxD$_{S}$ does not transfer iron to $_{H}$NifU.** Iron content of $_{H}$NifU (containing the core [$Fe_2S_2$] group), GrxD$_{S}$ (also purified with [$Fe_2S_2$] clusters), and $_{H}$NifU after being incubated and then separated from GrxD$_{S}$. Bars represent the average ± SD (n=3).

**Figure S8. Full length immunoblots shown in Figure 4.**

**Figure S9. R-NifU$_S$ does not transfer clusters to $_H$GrxD.** Iron content in reconstituted NifU (R-NifU$_S$), apo-$_H$GrxD, and apo-$_H$GrxD after being incubated with R-NifU$_S$ and separated with a Strep-tactin column. Bars represent the average ± SD (n=3).

**Figure S10. Apo-CA-$_H$NifU receives a [$Fe_2S_2$] cluster from GrxD$_S$ independent of reducing agent used.** Iron content of GrxD$_S$ and apo-CA-$_H$NifU prior to interaction, and in the flowthrough fraction (FT CA-$_H$NifU) obtained from passing through a Strep-tactin-column a solution in which apo-CA-$_H$NifU was incubated for 15 min with GrxD$_S$ in the presence of 5 mM GSH instead of DTT. Bars represent the average ± SD (n=3).

**Figure S11. Temperature dependent relaxation of X-EPR spectra.** A. X-band CW-EPR spectra of CA-$_H$NifU measured at different temperatures indicated in the figure. The signal intensity and line width are identical at 15 K and 50 K, suggesting that the S = ½ species is most likely a $[2Fe2S]^+$ cluster. B. X-band CW-EPR spectra of CA-$_H$NifU (A), GrxD$_S$ (B), flowthrough fraction after 15 min interaction of apo-CA-$_H$NifU and GrxD$_S$ (C) apo-CA-$_H$NifU (D) measured at 15 K. For B and D, the signals were multiplied by 20 times. A single S = ½ species with g = [2.02, 1.93, 1.89] is observed. The spin concentration of this species is 0.5 mM (A), 0 mM (B), 0.73 mM (C), and <10 µM (D).
