## Supplemental for "*Azotobacter vinelandii* glutaredoxin D delivers the core [Fe_2_S_2_] cluster to nitrogenase cofactor scaffold protein NifU": Supplemental Figures.pdf

**FIGURE S1**

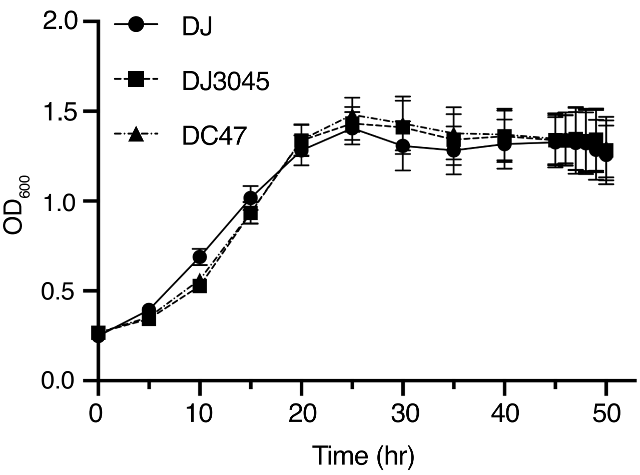

**FIGURE S2**

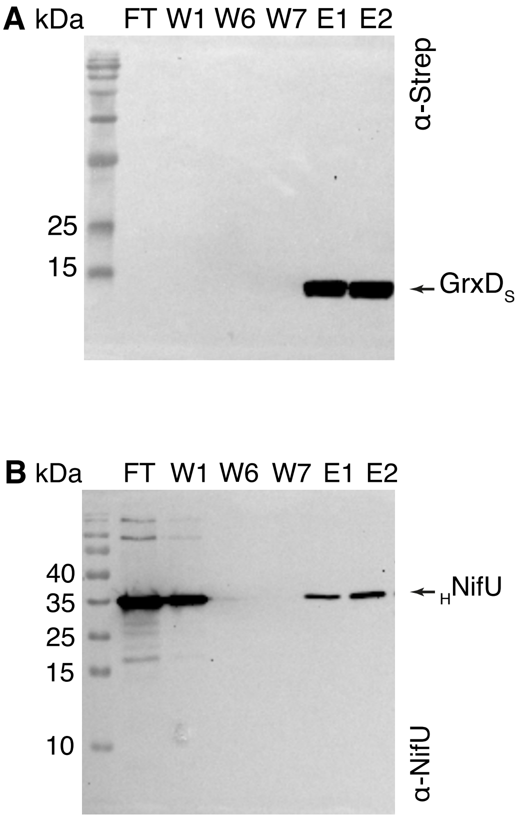

A kDa
FT W1 W6 W7 E1 E2
α-Strep
25
15
GrxD$_S$
B kDa
FT W1 W6 W7 E1 E2
40
35
25
15
10
$_H$NifU
α-NifU

**FIGURE S3**

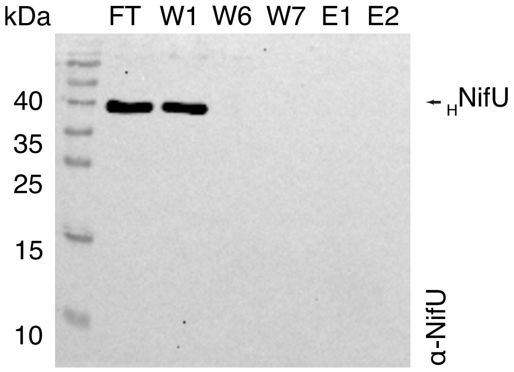

**FIGURE S4**

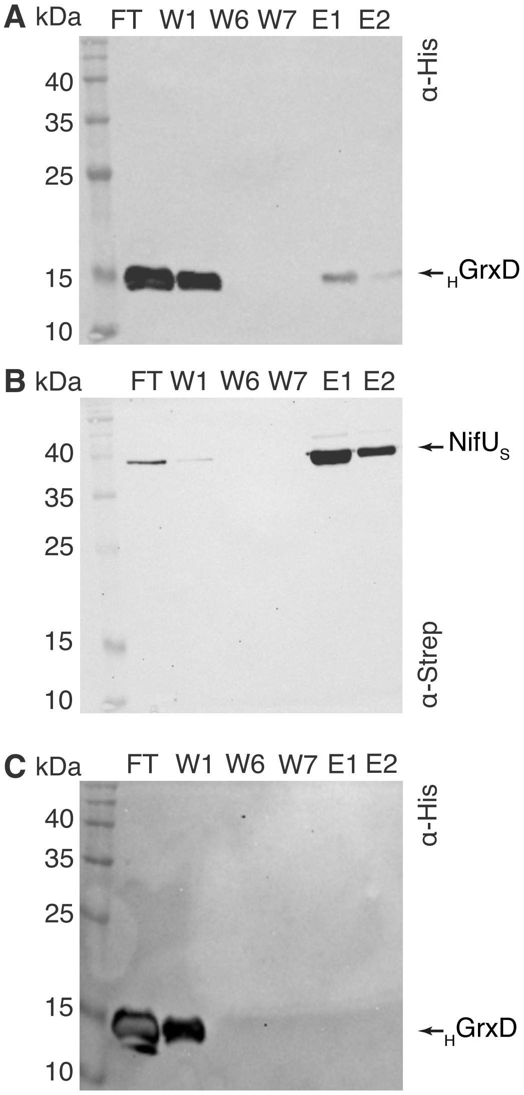
A kDa
FT W1 W6 W7 E1 E2
40
35
25
15
10
α-His
←HGrxD
B kDa
FT W1 W6 W7 E1 E2
40
35
25
15
10
←NifUS
α-Strep
C kDa
FT W1 W6 W7 E1 E2
40
35
25
15
10
α-His
←HGrxD

### FIGURE S5

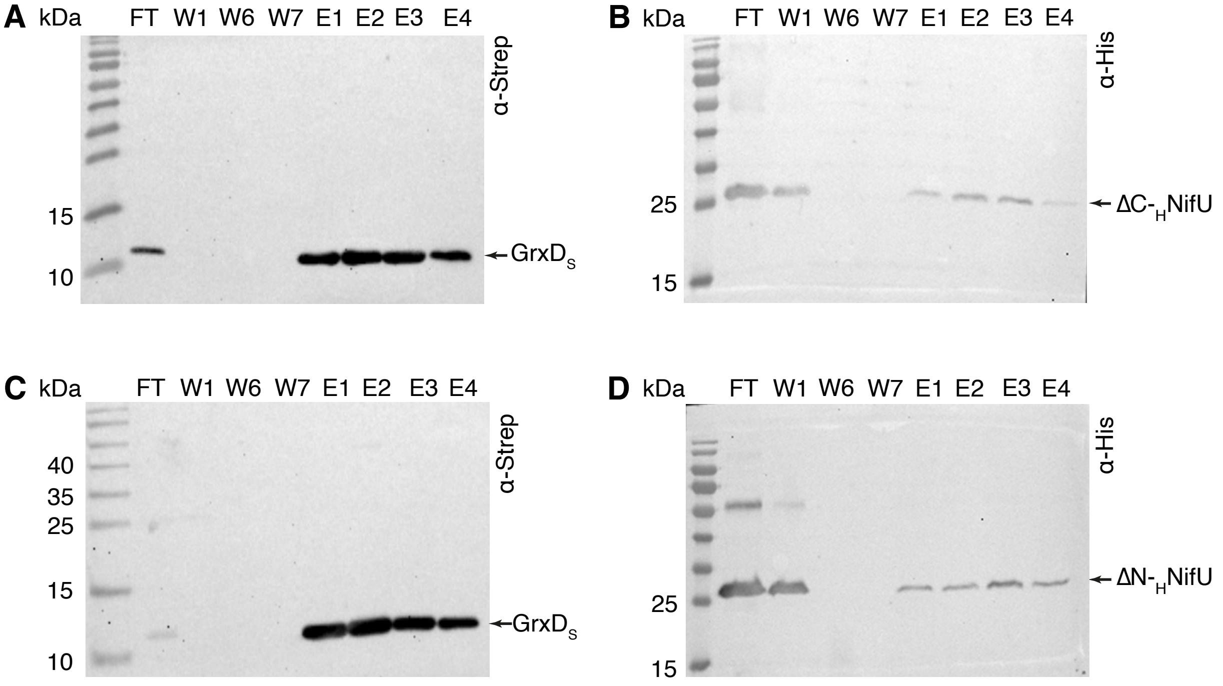
A
kDa
FT W1 W6 W7 E1 E2 E3 E4
α-Strep
15
10
GrxD$_S$
B
kDa
FT W1 W6 W7 E1 E2 E3 E4
α-His
25
15
ΔC-$_H$NifU
C
kDa
FT W1 W6 W7 E1 E2 E3 E4
α-Strep
40
35
25
15
10
GrxD$_S$
D
kDa
FT W1 W6 W7 E1 E2 E3 E4
α-His
25
15
ΔN-$_H$NifU

### FIGURA S6

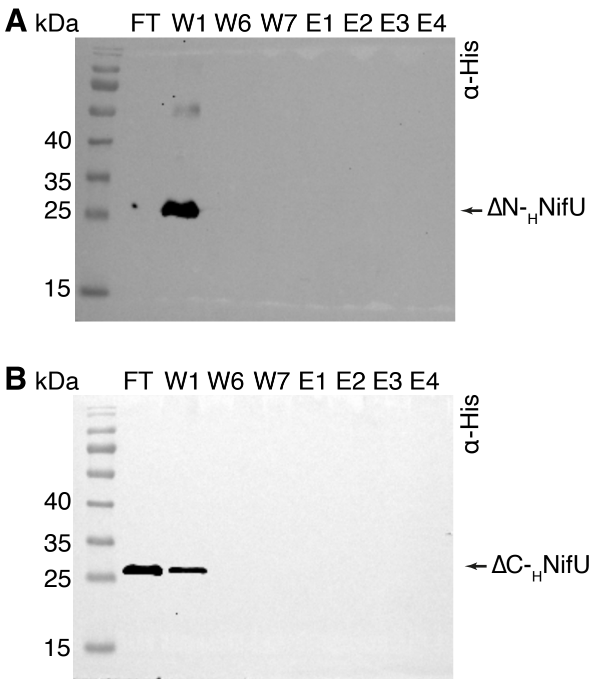

A kDa
FT W1 W6 W7 E1 E2 E3 E4
α-His
40
35
25
15
ΔN-$_{H}$NifU
B kDa
FT W1 W6 W7 E1 E2 E3 E4
α-His
40
35
25
15
ΔC-$_{H}$NifU

**FIGURE S7**

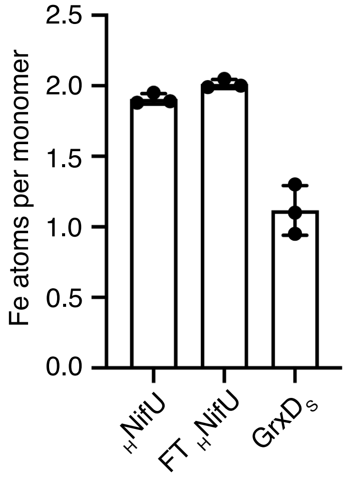

**FIGURE S8**

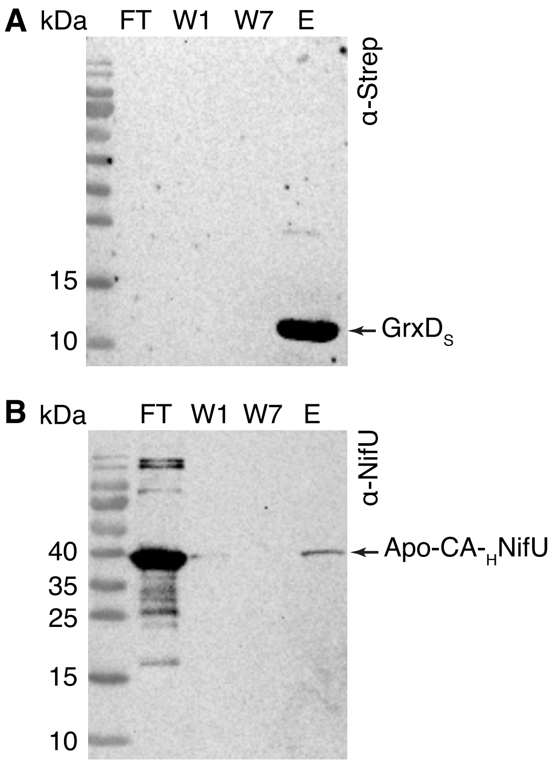
A kDa
FT W1 W7 E
α-Strep
15
10
GrxD$_S$
B kDa
FT W1 W7 E
α-NifU
40
35
25
15
10
Apo-CA-$_H$NifU

**FIGURE S9**

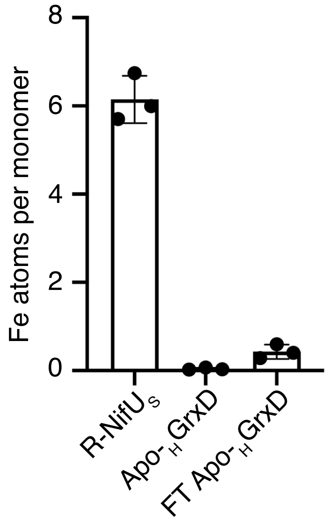

**FIGURE S10**

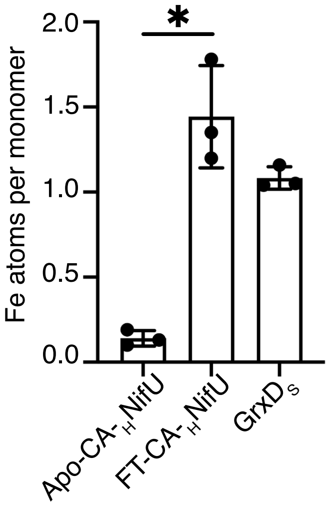

Fe atoms per monomer
2.0
1.5
1.0
0.5
0.0
*
Apo-CA-$_H$NifU
FT-CA-$_H$NifU
GrxD$_S$

#### FIGURE S11

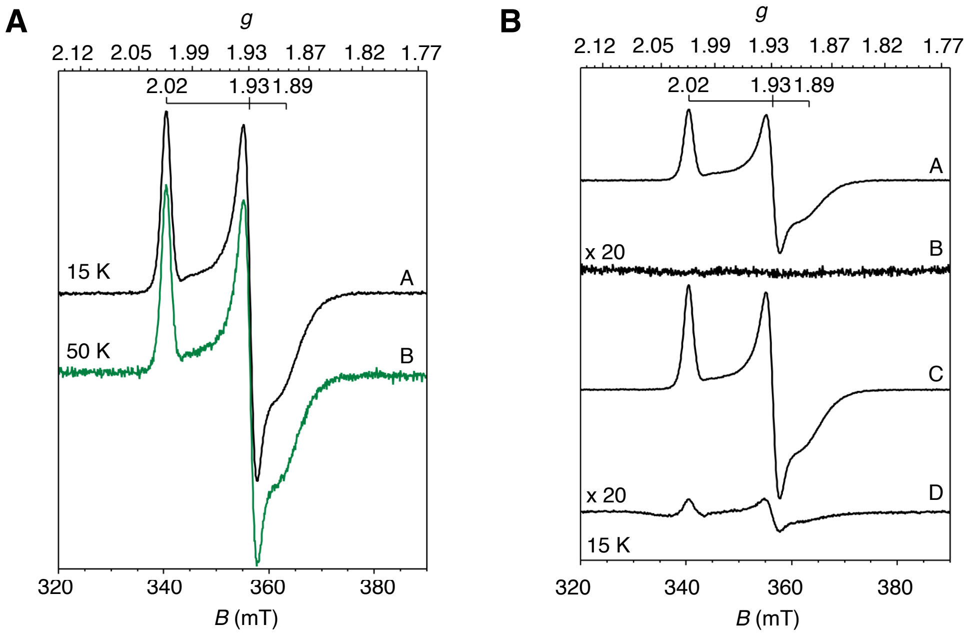
A
g
2.12
2.05
1.99
1.93
1.87
1.82
1.77
2.02
1.93
1.89
15 K
A
50 K
B
320
340
360
380
B (mT)
B
g
2.12
2.05
1.99
1.93
1.87
1.82
1.77
2.02
1.93
1.89
A
x 20
B
C
x 20
D
15 K
320
340
360
380
B (mT)
