## Supplemental for "*Azotobacter vinelandii* glutaredoxin D delivers the core [Fe_2_S_2_] cluster to nitrogenase cofactor scaffold protein NifU": Table S1.pdf

**Supplementary Table 1. Primers used in this study.**

| Use | Name | Sequence |
|---|---|---|
| $_{H}$GrxD cloning | 5*Nde*IGdxN-His | AGGCATATGATGGATATCATCGAAACCATT |
| | 3*Bam*HIGdxN-His | CCTGGATCCTCAAGCATCGGCTTTGTCGGC |
| GrxD$_{S}$ cloning | 5*Nco*IGdxC-Strep | AGGCCATGGATATCATCGAAACCATTAAAG |
| | 3*Nde*IGdxC-Strep | CCTCATATGAGCATCGGCTTTGTCGGCCGC |
| GrxD in frame mutant | FW-*Not*I-700UpGrx5 | AAAGCGGCCGCCAGGGATGCAGATCGTCGGAC |
| | 700DownGrx5*Eco*RI-RV | CCTGAATTCCAACCTCCAGGAGCTCGGCAA |
