## Supplemental for "*Azotobacter vinelandii* glutaredoxin D delivers the core [Fe_2_S_2_] cluster to nitrogenase cofactor scaffold protein NifU": Table S2.pdf

**Supplementary Table 2. Plasmid used in this work**

| Plasmid | Used for | Source |
|---|---|---|
| pRHB612 | To overexpress 9x His- *nifU* under T7 promoter, $Ap^R$ | Rubio laboratory collection |
| pDB303 | This plasmid carries the $Rif^R$ determinant | Dean laboratory collection |
| pDB1416 | To interrupts *scrB* gene with Gentamycin cartridge | Dean laboratory collection |
| pDB2123 | To overexpress streptag-*nifS* under T7 promoter, $Ap^R$. | Dean laboratory collection |
| pDB2174 | To overexpress *nifU*-streptag under T7 promoter, $Ap^R$ | Dean laboratory collection |
| pDB2678 | To generate a in frame deletion mutant *grxD* aa2-50 | Dean laboratory collection |
| pDB2679 | To restore *grxD* gene in a *grxD* mutant | Dean laboratory collection |
| pERN1 | To overexpress g*rxD*- strep tag under T7 promoter, $Ap^R$ | This work |
| pERN2 | To overexpress 9x His- g*rxD* under T7 promoter, $Ap^R$ | This work |
| pERN3 | To overexpress 9x His- ΔN - *nifU* under T7 promoter, $Ap^R$ | This work |
| pERN4 | To overexpress 9x His- ΔC- *nifU* under T7 promoter, $Ap^R$ | This work |
| pERN5 | To overexpress 9x His- CA- *nifU* under T7 promoter, $Ap^R$ | This work |
